# The *Plasmodium falciparum* cytoplasmic translation apparatus: a promising therapeutic target not yet exploited by clinically approved antimalarials

**DOI:** 10.1101/415513

**Authors:** Christine Moore Sheridan, Valentina E. Garcia, Vida Ahyong, Joseph L. DeRisi

## Abstract

The continued specter of resistance to existing antimalarials necessitates the pursuit of novel targets and mechanisms of action for drug development. One class of promising targets consists of the 80S ribosome and its associated components comprising the parasite translational apparatus. Development of translation-targeting therapeutics requires a greater understanding of protein synthesis and its regulation in the malaria parasite. Research in this area has been limited by the lack of appropriate experimental methods, particularly a direct measure of parasite translation. We have recently developed and optimized the PfIVT assay, an *in vitro* method directly measuring translation in whole-cell extracts from the malaria parasite *Plasmodium falciparum*.Here, we present an extensive pharmacologic assessment of the PfIVT assay using a wide range of known inhibitors, demonstrating its utility for studying activity of both ribosomal and non-ribosomal elements directly involved in translation. We further demonstrate the superiority of this assay over a historically utilized indirect measure of translation, S35-radiolabel incorporation. Additionally, we utilize the PfIVT assay to investigate a panel of clinically approved antimalarial drugs, many with unknown or unclear mechanisms of action, and show that none inhibit translation, reaffirming *Plasmodium* translation to be a viable alternative drug target. Within this set, we unambiguously find that mefloquine lacks translation inhibition activity, despite having been recently mischaracterized as a ribosomal inhibitor. This work exploits a direct and reproducible assay for measuring *P. falciparum* translation, demonstrating its value in the continued study of protein synthesis in malaria and its inhibition as a drug target.

**Author summary:** Novel antimalarial drugs are required to combat rising resistance to current therapies. The protein synthesis machinery of the malaria parasite *Plasmodium falciparum* is a promising unexploited target for antimalarial development, but its study has been hindered by use of indirect experimental methods which often produce misleading and inaccurate results. We have recently developed a direct method to investigate malaria protein synthesis utilizing whole-parasite extracts. In this work, we present an extensive characterization of the assay, using a panel of pharmacologic inhibitors with known mechanisms of action. We demonstrate the specificity of the assay in various stages of protein synthesis, as well as its improved accuracy and sensitivity in comparison to an indirect measure that has been the previous standard for the field. We further demonstrate that no current clinically available antimalarial drugs inhibit protein synthesis, emphasizing its potential as a target for drugs that will overcome existing resistance. Importantly, among the antimalarials tested was mefloquine, a widely used antimalarial that has recently been mischaracterized as an inhibitor protein synthesis. Our finding that mefloquine does not inhibit protein synthesis emphasizes the importance of using direct functional measurements when determining drug targets.

## Introduction

Despite ongoing efforts in its treatment and prevention, malaria remains a severe global health burden, with nearly half the world’s population at risk, and incidence of the disease actually increasing in the most recent years for which data are available [1]. Though malaria-related mortality has continued to decrease, the rise in incidence is particularly concerning in light of reduced investment worldwide in combatting malaria, combined with climate change and geopolitical instability that may contribute to a resurgence of the disease [1]. One compounding factor in the battle to eliminate malaria is the persistent emergence of drug resistance in the malaria parasite *Plasmodium falciparum* [1]. As combination therapies are the main defense against resistance, an important focus in therapeutic development is the identification of compounds with unique targets and novel mechanisms of action that are unlikely to be precluded by existing resistance mutations. Medicines for Malaria Venture (MMV) has recently demonstrated the potential of efforts directed at novel targets; two drugs currently showing great promise in clinical trials, SJ733 and cipargamin, inhibit the *P. falciparum* cation ATPase PfATP4, constituting a new class of drug [2,3].

One promising avenue for development of a novel target class is the inhibition of the *P. falciparum* ribosome, as well as other components of the translational machinery responsible for protein synthesis. Translation inhibitors have exhibited great clinical success as potent antibiotics, and in fact, several, including doxycycline and azithromycin, have found additional application as antimalarials, as they target ribosomes within the malaria parasite’s mitochondria and apicoplast, leading to loss of function of these organelles [4–6]. Interestingly, the *P. falciparum* cytoplasmic ribosome appears to occupy an evolutionary middle ground between prokaryotic and eukaryotic, differentiating it sufficiently from human ribosomes to yield a useful therapeutic window [5]. Indeed, a potent and highly selective inhibitor of the *P. falciparum* ribosome, M5717 (previously DDD107498), is currently in first-in-human study, validating the potential of the *P. falciparum* translational apparatus as an effective target for antimalarial drugs of this class [7].

To facilitate the identification of translation inhibitors, we previously developed a *P. falciparum* whole-cell extract-based *in vitro* translation assay (PfIVT), and successfully applied the technique to detect small molecule inhibitors in the MMV Malaria Box [8]. More recently, it has been suggested that the widely used drug mefloquine may inhibit the 80S ribosome of *P. falciparum* [9]. In addition, many currently approved antimalarial compounds lack a definitive mechanism of action, raising the possibility that some of these clinical therapies act through inhibition of translation. Here, we aimed to clarify which compounds truly exhibit inhibitory activity against the *Plasmodium falciparum* 80S ribosome and the associated translational apparatus. To do so, we compared a panel of antimalarial drugs (both clinical and pre-clinical) with well-characterized inhibitors of translation and other defined control compounds in the PfIVT assay, as well as in the S35-radioabel incorporation assay, a historically utilized indirect measure of translation. Importantly, we found that none of the current clinical therapeutics inhibit translation, including mefloquine. Regardless, testing of tool compounds shows that the PfIVT assay is capable of identifying not only translation inhibitors that directly interact with the ribosome, but also inhibitors of other non-ribosomal components of the translational machinery, demonstrating the broad utility of the assay for identifying novel malaria therapeutics that target *P. falciparum* translation.

## Results

### Extract optimization and quality control for the *Plasmodium falciparum in vitro* translation assay

To ensure reproducible consistency and robustness of the *P. falciparum in vitro* translation (PfIVT) assay, we performed extensive validation of parasite extracts. A detailed, step-by-step protocol is presented in the supplement. Only those extracts surpassing a rigorous activity threshold were utilized for the PfIVT assay, and extracts from individual harvests meeting this criterion were combined to generate large pools for use across many assays. Ribosome activity is especially sensitive to divalent cations, in particular magnesium concentration [10]. Therefore, we measured the magnesium concentrations of each PfIVT extract, and then determined the optimal amount of magnesium required by each extract in order to achieve maximal activity. Post-harvest magnesium concentrations were typically less than 2mM, whereas the maximum translational activity corresponded to a final PfIVT reaction concentration of approximately 4mM magnesium (Figs 1A and 1B). Upon determining optimal magnesium conditions for each pool of extract, kinetic curves were generated with 15-minute increments to establish the ideal incubation time for the assay (Fig 1C). This was necessary, since assay kinetics varied between extracts. Note that separate kinetic curves must also be established for the particular reporter utilized (in this case, firefly luciferase). To maintain maximal sensitivity to inhibitors and linearity of the assay, we conducted PfIVT experiments at the time point corresponding to 75-80% of the saturation signal (Fig 1C).

**Fig 1.**
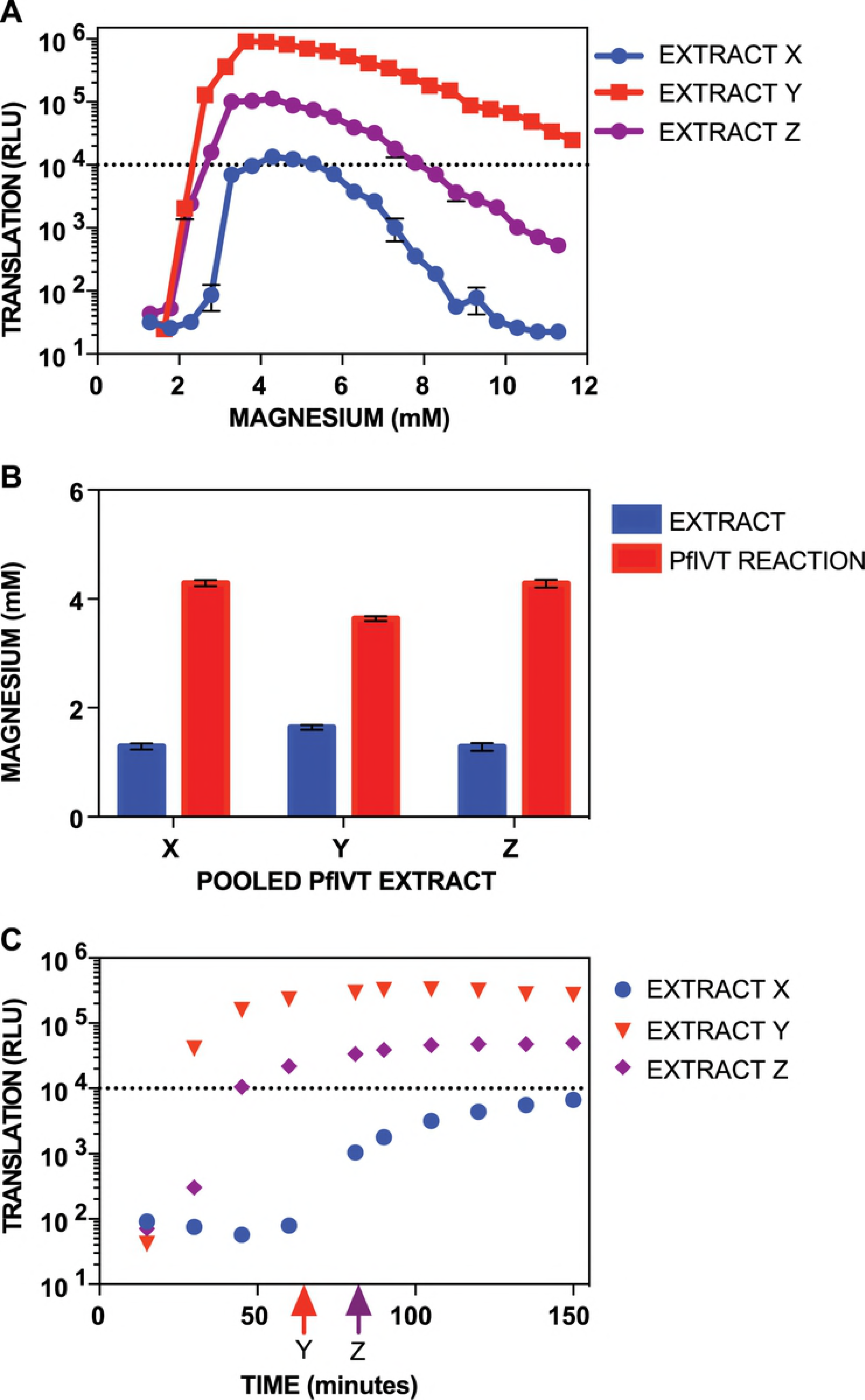
Optimization & quality control parameters of PfIVT extracts. A) Translational activity of 3 representative extracts (X, Y, and Z) over a range of reaction magnesium concentrations. B) Measured basal extract magnesium concentration (blue bars) and optimum translation reaction magnesium concentration (red bars) for each of 3 representative PfIVT extracts (X, Y, and Z). C) Kinetic curves for translational activity of each of 3 representative PfIVT extracts (X, Y, and Z) at the optimum reaction magnesium concentration shown in part B. Arrows indicate the timepoint to use for inhibition assays in the extracts meeting the activity threshold, indicating ∼75% of saturation signal. The dashed line at 104 relative luciferase units (RLU) represents the cutoff for acceptable translational activity for the assay. Extract X does not consistently meet the 104 RLU activity threshold and would not be used for PfIVT assays.

### Probing different stages of translation in a *Plasmodium falciparum* cellular extract system using tool compounds

The process of translation may be binned into three main stages: initiation, elongation, and termination [11,12]. In eukaryotes, this process is carried out by the 80S ribosome, comprised of a small (40S) and large (60S) subunit [11,12]. To further validate the PfIVT assay and investigate its capacity to interrogate the entirety of the normal activity of the 80S ribosome (and thus identify drugs inhibiting all steps of the process of translation), an extensive panel of previously characterized translational inhibitors was tested, both in the PfIVT assay, as well as in the historically utilized S35-radiolabelled amino acid incorporation assay. In contrast to the PfIVT assay, which directly measures activity of the 80S ribosome and the associated translational apparatus, S35 incorporation is an indirect measure of translation. Despite this, and the S35 incorporation assay’s resulting sensitivity to changes to upstream and parallel pathways, which often generating ambiguous or misleading results, it has remained a commonly used assay for studying parasite translation for lack of a better alternative [9].

Commercially available compounds that directly interact with the eukaryotic ribosome to inhibit translation initiation and/or elongation via a variety of mechanisms and binding sites, as well as several inhibitors of translation known to act upon non-ribosomal components of the translational machinery were tested (Tables 1 & 2). The eukaryote-specific inhibitors bruceantin and verrucarin A inhibit translation initiation through binding of mutually exclusive sites [12–15]. Suramin, also a specific inhibitor of the eukaryotic ribosome, inhibits both initiation and elongation through binding of multiple sites on the 40S, 60S and 80S ribosomes [16]. The eukaryote-specific elongation inhibitors tested are also distinct in their activities: cycloheximide and lactimidomycin overlap in their binding of the ribosome A-site, but differences in size and side-chains yield unique effects; anisomycin also overlaps cycloheximide’s binding site, but the two drugs bind the ribosome in distinct rotational conformations at different steps of elongation; homoharringtonine binds the A-site, but specifically inhibits re-initiating ribosomes; and nagilactone C inhibits both eEF-1α-dependent aminoacyl-tRNA loading and peptidyl transferase activity [12,13,17–19]. Halofuginone, also a specific inhibitor of eukaryote translation, does not interact with the ribosome, but instead inhibits glutamyl-prolyl-tRNA synthetase [20]. Puromycin was the sole pan-inhibitor tested, and acts as a tRNA mimetic that is incorporated into the nascent polypeptide chain, leading to its premature termination [21,22]. Negative controls were thiostrepton, a specific inhibitor of prokaryotic translation initiation and elongation; actinomycin D, an inhibitor of RNA polymerase II; tubercidin, an adenosine mimetic; and thapsigargin, a sarco/endoplasmic reticulum ATPase (SERCA) inhibitor [23–32].

**Table 1.** Mechanism of action and species specificity of translation inhibitors.

| COMPOUND | SPECIFICITY | MECHANISM OF ACTION | REFERENCE |
| --- | --- | --- | --- |
| bruceantin | eukaryotic (*Pf* cytoplasmic ribosomes) | inhibits initiation | [15] |
| verrucarin A | eukaryotic (*Pf* cytoplasmic ribosomes) | inhibits initiation, binds between P- & A-sites | [12–14] |
| suramin | eukaryotic (*Pf* cytoplasmic ribosomes) | inhibits initiation & elongation, multiple binding sites | [16] |
| anisomycin | eukaryotic (*Pf* cytoplasmic ribosomes) | inhibits elongation, binds A-site | [12,17] |
| cycloheximide | eukaryotic (*Pf* cytoplasmic ribosomes) | inhibits elongation, binds E-site | [12] |
| homoharringtonine | eukaryotic (*Pf* cytoplasmic ribosomes) | inhibits elongation on re-initiating ribosomes, binds A-site | [12,18] |
| lactimidomycin | eukaryotic (*Pf* cytoplasmic ribosomes) | inhibits elongation, binds E-site | [12] |
| nagilactone C | eukaryotic (*Pf* cytoplasmic ribosomes) | inhibits elongation, binds A-site | [12,19] |
| puromycin | pan-inhibitor | tRNA mimetic | [21,22] |
| halofuginone | eukaryotic | inhibits glutamyl-prolyl-tRNA synthetase | [20] |
| thiostrepton | prokaryotic (*Pf* mitochondrial & apicoplast ribosomes) | inhibits initiation & elongation | [23–29] |

**Table 2.** Mechanism of action of non-translation inhibitors.

| COMPOUND | MECHANISM OF ACTION | REFERENCE |
| --- | --- | --- |
| actinomycin D | RNA polymerase II inhibitor | [30] |
| tubercidin | adenosine mimetic | [31] |
| thapsigargin | SERCA inhibitor | [32–34] |

After determining the EC50 of each drug for the *P. falciparum* W2 strain in a 72-hour parasite growth assay, the drugs were characterized in both the S35 incorporation and PfIVT assays (Table 3)(Figs 2 and 3). Drugs were tested in the S35 and PfIVT assays at 0.1-, 1-, 10-, and 100-fold their determined growth assay EC50 in W2 parasites, except in cases where the highest concentration was constrained by solubility or available stock solution. The translation initiation inhibitors bruceantin and verrucarin A were both potent (nanomolar) inhibitors of S35 incorporation and PfIVT (Fig 2). All translation elongation inhibitors (anisomycin, cycloheximide, homoharringtonine, lactimidomycin, and nagilactone C) also strongly inhibited both S35 incorporation and PfIVT (Fig 2). Cycloheximide was additionally tested at 1000-fold its EC50, as it did not inhibit S35 incorporation at the lower concentrations tested, but did at this higher concentration, in line with inhibitory concentrations in recent reports, which also show that significantly higher concentrations of cycloheximide are required for complete, measurable inhibition of translation than for rapid and total parasite killing *in vivo* (S1 Fig) [8,9,35]. Suramin, which has been shown to inhibit both translation initiation and elongation, robustly inhibited PfIVT, but not S35 incorporation, likely due to poor cell permeability and the short timeframe of the S35 assay (2 hour drug pre-incubation followed by 2 hour radiolabel incorporation) (Fig 2). The tRNA mimetic puromycin, which induces premature termination of nascent polypeptides, inhibited both S35 incorporation and PfIVT with similar efficacy (Fig 2). Elucidating an even greater range of utility for the PfIVT assay, we found it to be capable of identifying inhibitors of non-ribosomal components of translation. The glutamyl-prolyl-tRNA synthetase inhibitor halofuginone inhibits both S35 incorporation and the PfIVT assay (Fig 2). In sum, these data demonstrate the ability of the PfIVT assay to interrogate both direct ribosomal activity, as well as extra-ribosomal components of the translational machinery.

**Table 3.** Half-maximal effective concentrations (nM) determined in *P. falciparum* growth inhibition assay.

| TEST COMPOUNDS | 72H GROWTH INHIBITION<br>EC50 (nM) | ANTIMALARIALS | 72H GROWTH INHIBITION<br>EC50 (nM) |
| --- | --- | --- | --- |
| bruceantin | 4.2 | MMV008270 | 2400 |
| verrucarin A | 0.6 | SJ733 | 60 |
| suramin | 1819 | M5717 (DDD107498) | 1 |
| anisomycin | 39 | quinine | 370 |
| cycloheximide | 0.6 | chloroquine | 333 |
| homoharringtonine | 6.8 | mefloquine | 25 |
| lactimidomycin | 22 | piperaquine | 26 |
| nagilactone C | 1310 | primaquine | 1849 |
| puromycin | 52 | monodesethyl amodiaquine<sup>a</sup> | 61 |
| halofuginone | 2 | lumefantrine | 4 |
| thiostrepton | 942 | dihydroartemisinin | 3 |
| actinomycin D | 10 |  |  |
| tubercidin | 168 |  |  |
| thapsigargin | 2900 |  |  |

<sup>a</sup> Active metabolite of amodiaquine

**Fig 2.**
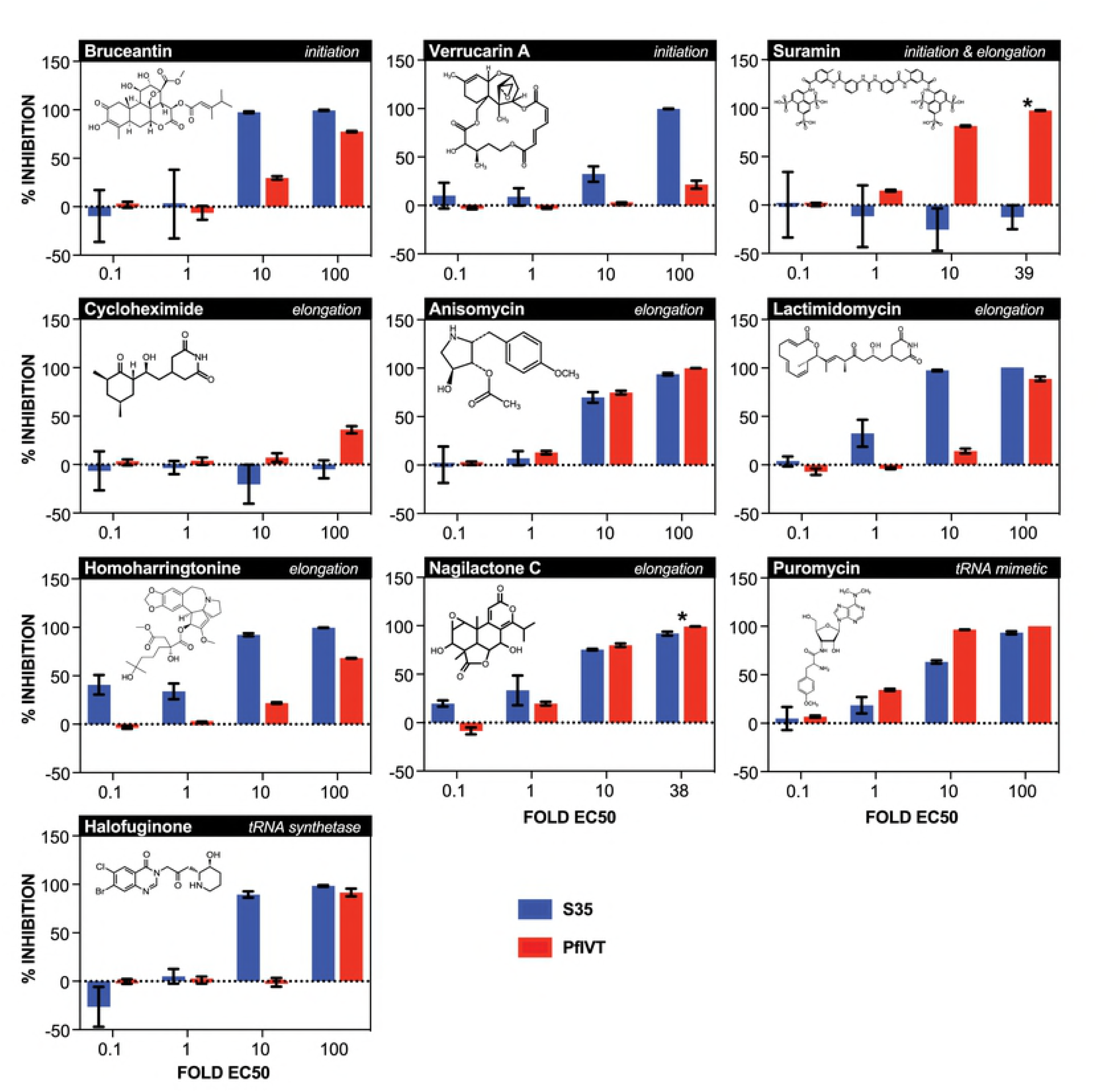
Dose-dependent inhibition of S35 incorporation and PfIVT assays by eukaryotic translation inhibitors. Dose-dependent inhibition, calculated as % inhibition, of S35 incorporation (blue bars) and PfIVT assays (red bars). Name of compound, mechanism of action, and molecular structure are displayed at top of each graph. Compounds were tested at 0.1-, 1-, 10-, and 100-fold the EC50 calculated in *in vivo* growth inhibition assay, except where upper concentration was limited by solubility, indicated by *.

**Fig 3.**
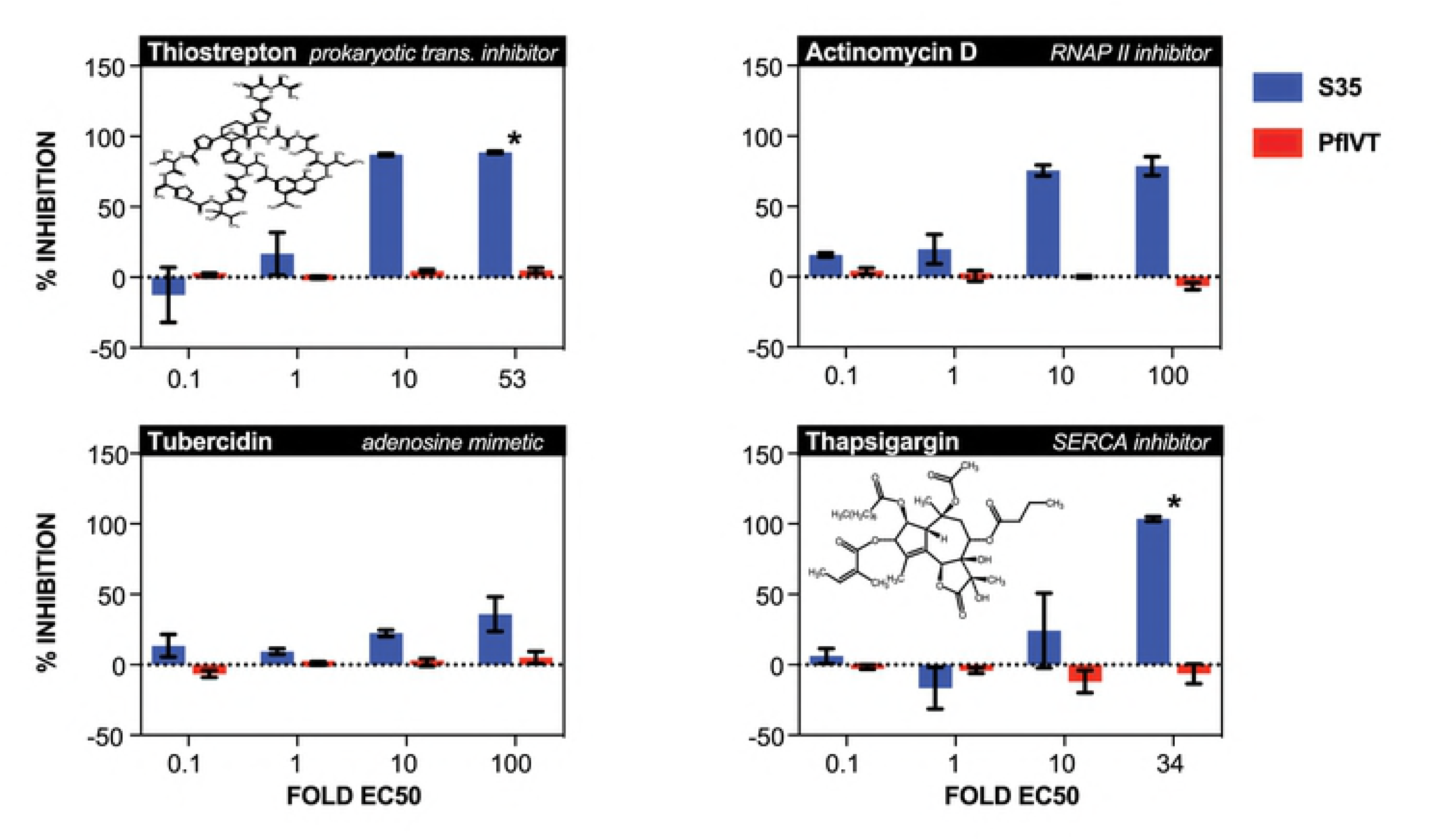
Dose-dependent inhibition of S35 incorporation and PfIVT assays by negative control compounds. Dose-dependent inhibition, calculated as % inhibition, of S35 incorporation (blue bars) and PfIVT assays (red bars) by negative control compounds: prokaryotic translation inhibitor and inhibitors of other (non-translation) cellular processes. Name of compound, mechanism of action, and molecular structure are displayed at top of each graph. Compounds were tested at 0.1-, 1-, 10-, and 100-fold the calculated EC50 calculated in *in vivo* growth inhibition assay, except where upper concentration was limited by solubility, indicated by *.

### The *Plasmodium falciparum in vitro* translation assay measures activity of cytoplasmic ribosomes

Importantly, all of the eukaryotic ribosome-specific inhibitors, which therefore should inhibit only *P. falciparum* cytoplasmic and not apicoplast or mitochondrial ribosomes, displayed inhibition in the PfIVT assay, with several achieving complete or near complete blocking of translation (suramin, anisomycin, lactimidomycin, nagilactone C) (Fig 2). In addition, the prokaryotic ribosome-specific inhibitor thiostrepton did not inhibit the PfIVT assay at any concentration tested, despite inhibiting the S35 assay at concentrations above its determined EC50 (Fig 3). Thiostrepton is known to have multiple targets apart from ribosomes in eukaryotes and has been shown to induce an ER stress response with a phenotype similar to thapsigargin, which likely accounts for its activity in the S35 assay [36–38].

### The S35 incorporation assay is not a reliable indicator of direct translation inhibition

Although it is well documented in other model systems (i.e. yeast) that the S35-radiolabeled amino acid incorporation assay is an indirect measure of translation and can, as such, generate many misleading artifacts, this has not yet been characterized carefully with respect to *Plasmodium spp.* [33,34]. Despite this, several studies in *Plasmodium* have relied on this indirect measure as a primary readout of translation [9,39]. To address this and further determine the specificity of the PfIVT assay relative to the S35 uptake assay, we tested a panel of small molecules that are known to inhibit cellular processes other than translation (Tables 2 and 3). Not surprisingly, actinomycin D, an inhibitor of transcription targeting RNA Polymerase II, and the SERCA inhibitor thapsigargin both exhibited strong inhibition in the S35 incorporation assay, but had no effect in the PfIVT assay (Fig 3). Tubercidin, an adenosine mimetic, had a modest inhibitory effect on S35 incorporation, but, again, negligible effect in the PfIVT assay (Fig 3). These data confirm that the PfIVT assay directly measures translation, and highlight the lack of translation specificity of the S35 incorporation assay.

### Analysis of clinically-approved antimalarials reveals that none, including mefloquine, inhibit the 80S ribosome

We next sought to test a panel of clinically approved antimalarial drugs with undefined or disputed mechanisms of action, to determine whether any might act through direct inhibition of translation, subjecting these drugs to the same battery of assays described above (*P. falciparum* growth, PfIVT, and S-35 incorporation) (Tables 3 and 4). Chloroquine and piperaquine were mild inhibitors of the S35 incorporation assay at the highest drug concentrations tested (Fig 4). Quinine, lumefantrine, primaquine, monodesethyl amodiaquine (the active metabolite of amodiaquine), and dihydroartemisinin were moderate-to-strong inhibitors of the S35 incorporation assay (Fig 4). SJ733, an inhibitor of the sodium transporter PfATP4, and a clinical candidate currently in Phase I trials, exhibited strong inhibition in the S35 incorporation assay (Fig 4). Notably, none of these antimalarial drugs inhibited the PfIVT assay. However, primaquine cannot be ruled out with complete certainty as an inhibitor of translation, as its active metabolite may not be produced in an *in vitro* setting, and it does show moderate activity in the S35 incorporation assay [40].

**Fig 4.**
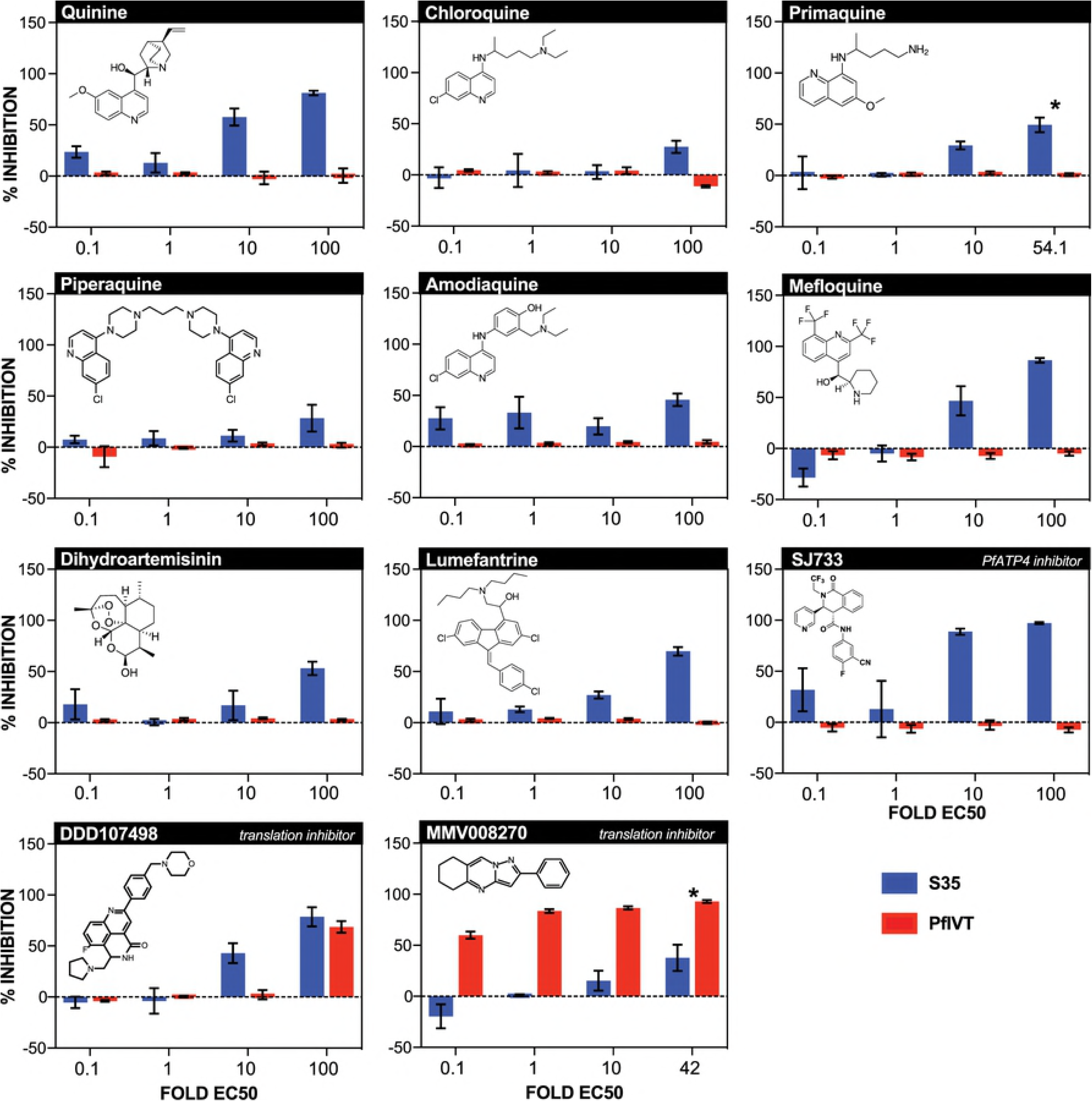
Dose-dependent inhibition of S35 incorporation and PfIVT assays by antimalarial compounds. Dose-dependent inhibition, calculated as % inhibition, of S35 incorporation (blue bars) and PfIVT assays (red bars) by pre-clinical and clinically-approved antimalarial compounds. Name of compound, mechanism of action (where definitively known), and molecular structure are displayed at top of each graph. Compounds were tested at 0.1-, 1-, 10-, and 100-fold the calculated EC50 calculated in *in vivo* growth inhibition assay, except where upper concentration was limited by solubility, indicated by *.

We also included several drugs (clinical and pre-clinical) that have recently been reported to inhibit translation (Tables 3 and 4)[7–9]. MMV008270 was a moderate inhibitor of the S35 incorporation assay, while mefloquine and DDD107498 robustly inhibited S35 incorporation (Fig 4). Strikingly, while DDD107498 and MMV008270 inhibited the PfIVT assay, mefloquine failed to do so (Fig 4). Interestingly, MMV008270 was an exceptionally effective inhibitor of translation in the PfIVT assay at all concentrations tested, significantly outperforming the S35 incorporation assay (Fig 4). These data reveal that mefloquine has recently been mischaracterized as a ribosome inhibitor through use of the S35 incorporation assay, when it does not, in fact, directly inhibit translation [9].

To further validate the PfIVT data regarding mefloquine, we repeated the PfIVT assay, alongside a commercially available rabbit reticulocyte *in vitro* translation assay (RRIVT), with a full titration of drug to determine half maximal effective values for both mefloquine and DDD107498 (Fig 5). As expected, the positive control cycloheximide was a robust inhibitor of both translation systems (PfIVT IC50: 31.91nM, RRIVT IC50: 37.8nM), while DDD107498 was a potent inhibitor of *P. falciparum*, but not rabbit reticulocyte translation, confirming the reported high *P. falciparum* selectivity of DDD107498 (PfIVT IC50: 60.5nM)(Fig 5).In contrast, mefloquine failed to inhibit in either the PfIVT or RRIVT assay, even at concentrations as high as 20uM (Fig 5). The reported binding site of mefloquine to the 80S ribosome is on the highly conserved ribosomal protein uL13; if this were indeed the active binding site of the drug, mefloquine should inhibit the RRIVT assay [41]. To rule out the possibility that mefloquine solubility may be a confounding factor in the IVT assays, completed PfIVT reactions with a dilution series of mefloquine or DMSO control were centrifuged at high speed, sterile-filtered, and the resulting supernatant was used as the input for an *in vivo* growth assay. EC50 values were comparable between the IVT reaction supernatant containing mefloquine (12.31nM) and mefloquine alone (4.17nM), thus demonstrating that mefloquine is soluble in the PfIVT assay (S2 Fig). These data make clear that mefloquine does not act through inhibition of the *P. falciparum* ribosome, nor through other direct inhibition of the translational machinery.

**Fig 5.**
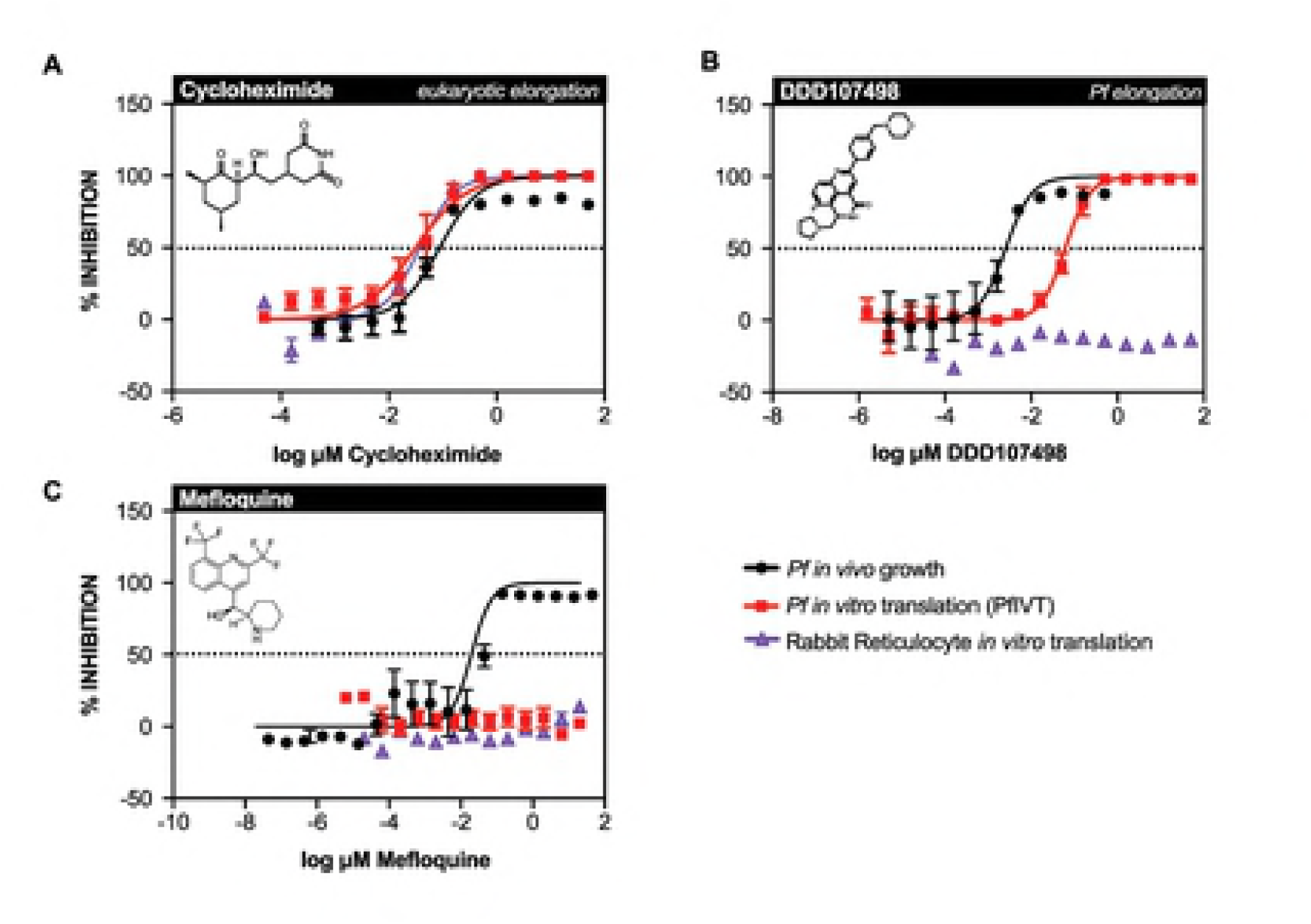
Dose-response curves of Pf growth, PfIVT, and RRIVT for mefloquine and controls. Dose-response curves comparing inhibition, calculated as % inhibition, of *P. falciparum* growth assay (black), *P. falciparum in vitro* translation assay (red), and commercially available rabbit reticulocyte *in vitro* translation assay (purple). Name of compound, mechanism of action (where definitively known), and molecular structure are displayed at top of each graph.

## Discussion

This work presents an extensive dissection and validation of the whole-cell extract-derived PfIVT assay, the only reported direct measure of *P. falciparum* translation to date. Through probing the assay with numerous small molecule inhibitors of translation, exhibiting a diversity of binding sites and mechanisms of action, as well as a variety of well-characterized tool compounds inhibiting non-translational pathways, we demonstrate that the PfIVT assay specifically measures *P. falciparum* cytoplasmic ribosome activity. *In vitro* translation extracts are inherently difficult to make, and even more so for an intraerythrocytic parasite. However, when subjected to stringent quality control and careful optimization, the PfIVT assay reliably and specifically identifies inhibitors of translation initiation and elongation, as well as inhibitors of non-ribosomal proteins necessary for translation, such as tRNA synthetase.

The PfIVT assay is particularly valuable to the study of *P. falciparum* translation as a direct measure of translation, as opposed to the indirect measures to which the field has historically been constrained, such as incorporation of radiolabeled amino acids *in vivo*. Importantly, our data show the PfIVT assay is significantly more specific, and in some cases more sensitive, than S35-radiolabel incorporation in identifying small molecule inhibitors of translation. Indeed, the PfIVT assay specifically identified all eukaryotic translation inhibitors tested, while S35-radiolabel incorporation was prone to false-positives. We found that none of the clinically approved antimalarials tested are inhibitors of translation, emphasizing the potential for translation as a useful therapeutic target, as there is unlikely to be pre-existing mechanism-specific resistance to any identified candidates resulting from use of these drugs. It is notable that mefloquine, in contrast to other previously reported translation inhibitors, did not exhibit any inhibitory activity. Mefloquine was likely mischaracterized as an 80S ribosome inhibitor through a combination of non-specific inhibition of S35 incorporation, as well as artifacts arising from cryo-EM structures obtained under the non-physiologic condition of 10mM magnesium – well above the ∼4mM magnesium that we have found to be optimal for translation (Fig 1)[9].

While the PfIVT assay exhibits clear benefits over existing methodologies for the study of *P. falciparum* translation, we acknowledge that the technique has several limitations. As is the case with *in vitro* translation systems in other organisms, the current assay is likely biased toward the study of non-cap-dependent initiation and elongation. We utilized uncapped mRNA in this study to focus specifically on activity of the 80S ribosome itself, rather than the cap-recognition apparatus. It is possible that utilization of capped mRNA in future studies would facilitate interrogation of cap-dependent translation initiation. Likewise, there are few characterized pharmacologic inhibitors of eukaryotic translation termination, none of which are currently commercially available; thus, the PfIVT system, as described, may not be sensitive to all specific inhibitors of translation termination. Additionally, some translation inhibitors, such as homoharringtonine, demonstrated greater potency in the S35 incorporation and growth inhibition assays than in PfIVT. Such variation between the two assays may suggest off-target effects of these drugs, or differences between whole living cells and cellular extracts.

Determining the true molecular targets of antimalarials is critical to improved therapeutic development. Exploiting differences between *P. falciparum* and mammalian ribosomes remains a promising avenue, as evidenced by the potent and discriminating drug DDD107498. Here, we have shown that orthogonal biochemical assays may be used to test hypotheses generated by structural data and cell-based assessments. Our investigation of mefloquine reaffirms that direct functional measurements of drug activity are critical to identifying the genuine molecular targets of drugs. Importantly, we show that the PfIVT assay is a uniquely direct measure of *P. falciparum* translation that can be used to facilitate a better understanding of the specifics of *P. falciparum* protein synthesis, with potentially great consequences for antimalarial therapeutic development.

## Methods

### Drug stocks

*In vivo* growth and *in vitro* translation measurements were performed using the same drug dilutions. The antimalarial drugs chloroquine, dihydroartemisinin, lumefantrine, monodesethyl amodiaquine, piperaquine, primaquine, and quinine were a generous gift from Dr. Phil Rosenthal of UCSF. SJ733 was generously provided by Dr. Kip Guy of St. Jude Children’s Research Hospital. All other compounds were purchased from the indicated vendors: DDD107498 (Apexbio #A8711-5), mefloquine hydrochloride (Sigma-Aldrich #M2319), emetine (Sigma-Aldrich #E2375), cycloheximide (Fisher #AC35742-0010), MMV008270 (Vitas-M Laboratory #STK591252), actinomycin D (Sigma-Aldrich #A1410), tubercidin (Sigma-Aldrich #T0642), thapsigargin (Sigma-Aldrich #SML1845), ionomycin (Sigma-Aldrich #407951), thiostrepton (Sigma-Aldrich #598226), bruceantin (Toronto Research Chemicals #B689310), verrucarin A (Sigma-Aldrich #V4877), anisomycin (Sigma-Aldrich #A5862), homoharringtonine (Sigma-Aldrich #SML1091), lactimidomycin (EMD Millipore #506291), nagilactone C (BOC Sciences #24338-53-2), suramin sodium salt (Sigma-Aldrich #S2671), puromycin (Thermo Fisher #A1113803), halofuginone (Sigma-Aldrich #32481.

### *Plasmodium falciparum* strain and culturing

*Plasmodium falciparum* W2 (MRA-157) was obtained from MR4. Parasites were grown in human erythrocytes (2% hematocrit) in RPMIc (RPMI 1640 media supplemented with 0.25% Albumax II (GIBCO Life Technologies), 2 g/L sodium bicarbonate, 0.1 mM hypoxanthine, 25 mM HEPES (pH 7.4), and 50 μg/L gentamicin), at 37 °C, 5% O2, and 5% CO2. Cells were synchronized with 5% sorbitol treatment for two generations to achieve high synchronicity.

### Growth inhibition assays

2μL of serial drug dilutions in 100% DMSO were dispensed in triplicate to 96-well plates utilizing the LabCyte ECHO acoustic liquid handler. 198µL of *P. falciparum* W2 cultures were added. Growth was initiated with ring-stage parasites at 0.8% parasitemia and 0.5% hematocrit. Plates were incubated at 37 °C, 5% O2, and 5% CO2 for 72 hours. Growth was terminated by fixation with 1% formaldehyde, and parasitized cells were stained with 50nM YOYO-1 (Invitrogen). Parasitemia was determined by flow cytometry on the BD LSRII, analyzed using FlowJo software version 10, and EC50 were curves plotted by GraphPad Prism. Two biological replicates were performed for each drug.

### Generation and quality control of extracts for *Plasmodium falciparum in vitro* translation assay

For PfIVT harvests, one liter of synchronized parasite culture in 2-4% hematocrit was grown in two 500mL HYPERFlask M vessels (Corning), and media was changed every 8-12 hours, with the final media change at 4-8 hours prior to harvest. Parasites were harvested in the late trophozoite stage at 15-20% parasitemia by centrifugation for 5 min at 1500×*g* at room temperature, followed by removal of media and addition of ice-cold 0.025-0.05% final saponin in Buffer A (20 mM HEPES pH 8.0, 2 mM Mg(OAc)2, 120 mM KOAc). Due to variations between and within lots, saponin stocks were prepared in large volumes, aliquoted, and stored at −20°C. Percentage utilized for each batch of aliquots was determined empirically through pairwise testing of concentrations (1 for each HYPERFlask) and assessed via resulting activity of PfIVT extracts. Saponin-lysed pellets were centrifuged at 4°C and 10,000×*g* for 10 minutes and washed twice with ice-cold Buffer A. Supernatant was carefully removed, and washed pellets were resuspended in an equal volume of Buffer B2 (20 mM HEPES pH8.0, 100 mM KOAc, 0.75 mM Mg(OAC)2, 2 mM DTT, 20 % glycerol, 1X EDTA-free protease inhibitor cocktail (Roche)), flash frozen, and stored in −80°C freezer until the sample was ready to homogenize.

Frozen pellets were thawed on ice and added to a 3-mL Luer lock syringe, which was then secured onto a pre-chilled cell homogenizer containing a 4μm-clearance ball bearing (Isobiotec, Germany) that was pre-washed with ice-cold Buffer B2. Homogenate was passed between two syringes 20 times on ice, either by hand or by use of a custom robot built to accommodate the cell homogenizer (42). Lysate was immediately centrifuged at 4°C and 16,000×*g* for 10 minutes, and the supernatant (the resulting PfIVT extract) was transferred to a fresh tube, with a small (100μL) aliquot set aside for activity testing. Extracts and test aliquots were flash-frozen and stored at −80°C. Test aliquots from multiple harvests were thawed on ice and tested in batches in the PfIVT assay (see below) across a small range of magnesium concentrations with a 2 hour incubation time, using a firefly luciferase reporter. Extracts that surpass the activity threshold of 104 relative luciferase units (RLU) were then thawed on ice and combined to generate large volume pools. Extract pools were flash-frozen in 200μL aliquots and stored at −80°C. Extract pools were tested across a range of magnesium concentrations via PfIVT assay to determine the optimum magnesium concentration. Once magnesium concentration has been determined, pools are then tested in the PfIVT assay in 15 minute incubation time points up to 150 minutes to determine the kinetics of the extract pool, and thus the appropriate incubation time for the pool (∼75-80% of maximum signal, within the linear range of the extract’s kinetic curve). Kinetics must be separately assessed for each reporter used (i.e. if a nanoluciferase reporter is used instead of firefly luciferase).

### Magnesium concentration assays

Baseline magnesium levels of the PfIVT extracts were measured using a magnesium-dependent enzyme-based colorimetric assay kit (Sigma-Aldrich #MAK026). Two biological replicates of a dilution series of each extract were tested in duplicate with each of two separate kits, following the protocol provided with the kit. In brief, 10μL of each PfIVT extract (neat, or diluted 1:4 or 1:10 with ddH2O) added to 10μL ddH2O, along with a standard curve, was combined with 50μL of master reaction mix (35μL magnesium assay buffer, 10μL developer, 5μL magnesium enzyme mix), and incubated for 10 minutes with shaking at 37°C. 450nm absorbance was read immediately after the initial incubation, and every 5 minutes thereafter on a Tecan plate reader until the highest A450 approached (but did not exceed) 1.5X the initial reading. Values were fitted to, and interpolated from, the standard curve using Prism GraphPad.

### *Plasmodium falciparum in vitro* translation assay

*P. falciparum in vitro* translation (PfIVT) reactions were carried out in skirted v-bottom 96-well PCR plates (BioRad) and sealed with adhesive aluminum foil plate seals (Beckman Coulter, Indianapolis, IN, USA). 200nL of drug in 100% DMSO was dispensed in duplicate to appropriate wells of the plate utilizing a Labcyte ECHO acoustic liquid handler. 19.8μL of PfIVT reaction mix (per 20μL: 14μL extract, 1 μg T7-transcribed firefly luciferase mRNA, 10 µM amino acid mixture, 20 mM HEPES/KOH pH 8.0, 75 mM KOAc, 2 mM DTT, 0.5 mM ATP, 0.1 mM GTP, 20 mM creatine phosphate, 0.2 μg/μl creatine kinase, and the appropriate amount of Mg(OAc)2 as determined for the particular pool of extract) was then dispensed to each well using Rainin E4 12-channel electronic pipettes (Rainin Instruments, Oakland, CA, USA). Reactions were incubated at 37°C for the appropriate amount of time as determined for the particular pool of extract. After incubation, the reactions were placed on ice, then quenched through transfer to a 96-well LUMITRAC 200 flat-bottom white assay plate (Greiner Bio-One, Monroe, NC, USA) containing 2μL of 50μM cycloheximide (dispensed using the Labcyte ECHO), then immediately centrifuged to combine the PfIVT reaction with the cycloheximide for a final concentration of 5 µM cycloheximide. Reactions were assayed using the Promega GloMax-Multi + microplate reader with a three-second delay and three-second integration after addition of 200μL luciferin reagent dispensed at a speed of 200μL/second (firefly luciferin reagent: 20 mM Tricine, 2.67 mM MgSO4×7H2O, 0.1 mM EDTA, 33.3 mM DTT, 530 μM ATP, 270 μM Acetyl CoEnzyme A, 1 mM D-Luciferin, 265 μM Magnesium Carbonate Hydroxide, pH 8.15). Three biological replicates were performed in duplicate for each drug. IC50 curves were plotted by GraphPad Prism.

### Rabbit reticulocyte *in vitro* translation assay

Rabbit reticulocyte *in vitro* translation assays were performed as described in Ahyong *et al* with the exception that the final [DMSO] for MMV008270 was 2.5% and all other final [DMSO] = 0.55% [8]. IC50 curves were plotted by GraphPad Prism.

### S35 incorporation assay

#### Parasite purification

Synchronized parasites were cultured in 2% hematocrit at 10-15% parasitemia, and MACS purified at the late trophozoite stage to remove uninfected erythrocytes using standard protocols. In brief, at least two LD MACS Separation columns (Miltenyi Biotech) per 50mL of culture were washed with 1.25mL of pre-warmed RPMIc. Next, cultures were added to the columns 5mL at a time and allowed to gravity filter at 37°C. Finally, the columns were rinsed with 2.5mL of pre-warmed RPMIc, removed from the magnetic stand, and eluted with 2mL of pre-warmed RPMIc.

#### Drug treatment and S35 labeling

1μL of drug in 100% DMSO was dispensed to each well of a 96-well round-bottom culture plate utilizing the Labcyte ECHO acoustic liquid handler. 2×107 MACS-purified parasites in 199μL of RPMIc were then added to each well. Parasites were incubated with drug for 2 hours at 37°C, 5% CO2, 5% O2. Next, samples were transferred from 96-well plates to 1.5mL screw-cap microfuge tubes. 35μCi of EasyTag™ Express S35 Protein Labeling Mix (Perkin Elmer) diluted to 10μL with RPMIc was added to each tube. Reactions were incubated at 37°C with mild shaking for 2 hours.

#### Washing and lysis

After incubation, cells were pelleted and 160μL of supernatant was removed. Parasites were then washed with 200μL of ice-cold PBS containing 50μM cycloheximide four times. After the final wash, all supernatant was removed and samples were resuspended in 15μL of 2X SDS buffer (100mM Tris-Cl pH 6.8, 4% SDS, 20% glycerol, 0.1M DTT). Samples were boiled at 98°C for 5 minutes and stored at −20°C.

#### Scintillation counting

Samples were thawed at room temperature, boiled for 5 minutes at 98°C, and spun at max speed in a tabletop microcentrifuge for 10 minutes. 10μL of supernatant per sample was placed on a 0.45μm nitrocellulose membrane (HAWP02400 from Millipore). Each membrane was washed 4 times with 15mL of TBS-T then placed in a 20mL HDPE scintillation vial (Fisher Scientific) with 8mL of Ecoscint A (National Diagnostics). S35 counts were measured for 1 minute using a Beckmann coulter, LS 6500 Multi-purpose Scintillation Counter. Three biological replicates were performed for each drug.

### Mefloquine solubility assay

PfIVT extracts were incubated with a dilution series of mefloquine or DMSO control for 90 minutes. All PfIVT conditions were the same as above, except without addition of cycloheximide to stop translation. Reactions were centrifuged at 16,100x*g* for 10 minutes at room temperature; resulting supernatant was then filtered and added to cultures for the *P. falciparum* growth inhibition assays as described above.

## ACKNOWLEDGEMENTS

We thank Dr. Phil Rosenthal for providing stocks of several antimalarial drugs.

## Supporting information

**S1 Fig. Dose-dependent inhibition of S35 incorporation and PfIVT assays by cycloheximide.** Dose-dependent inhibition of S35 incorporation (blue bars) and PfIVT assays (red bars) by the translation inhibitor cycloheximide, tested up to 1000-fold (**) the EC50 calculated in *P. falciparum* growth inhibition assay.

**S2 Fig. Mefloquine solubility assay.** Dose-dependent inhibition of *P. falciparum in vivo* growth by mefloquine in PfIVT extract post-PfIVT reaction (Mefloquine IVT Extract), mefloquine alone (Mefloquine No Extract), or DMSO control in PfIVT extract post-PfIVT reaction (Extract + DMSO).

**S3 Fig. Flowchart of saponin batch calibration for erythrocyte lysis.** Saponin amounts for RBC lysis are empirically determined through pairwise comparison for each preparation/batch of saponin. 3 to 5 harvests and pairwise tests will be required to determine the ideal amount of saponin for a given batch. Volumes indicated on the flowchart are for the volume of 0.15% saponin (in Buffer A) to be added to parasites in Buffer A and to a total volume of 50mL. Harvest 1 should be tested with 8mL and 10mL saponin. Subsequent pairs for testing are determined by following the arrows on the flowchart: if 8mL yields the more active extract in Harvest 1, Harvest 2 will compare 8mL with 6mL saponin; if 6mL yields the more active extract in Harvest 2, Harvest 3 will compare 6mL with 7mL saponin, and so on, until a final value has been reached.

**S4 Fig. Flowchart for PfIVT extract quality control and pooling.** Individual harvests are first tested at different magnesium concentrations; those that achieve 104 RLU activity threshold are pooled. Pooled extract is aliquoted, and a test aliquot is utilized to first determine ideal magnesium concentration, then ideal incubation time. Remaining aliquots are utilized for PfIVT assays at the determined magnesium & kinetic conditions.

**S1 File. Method for preparation and calibration of saponin.**

**S2 File. Method for culturing and extract generation: step-by-step protocol.**

**S3 File. Method for PfIVT assay: step-by-step protocol.**

